# Multisite imaging of neural activity using a genetically encoded calcium sensor in the honey bee

**DOI:** 10.1101/2022.04.22.489138

**Authors:** Julie Carcaud, Marianne Otte, Bernd Grünewald, Albrecht Haase, Jean-Christophe Sandoz, Martin Beye

**Author notes:** Corresponding author: Julie Carcaud, Evolution, Genomes, Behavior and Ecology, Université Paris-Saclay, CNRS, IRD, Gif-sur-Yvette, France. These authors jointly supervised this work (Albrecht Haase, Jean-Christophe Sandoz, Martin Beye).

## Abstract

Understanding of the neural bases for complex behaviors in Hymenoptera insect species has been limited by a lack of tools that allow measuring neuronal activity simultaneously in different brain regions. Here, we developed the first pan-neuronal genetic driver in a Hymenopteran model organism, the honey bee, and expressed the calcium indicator GCaMP6f under the control of the honey bee *synapsin* promoter. We show that GCaMP6f is widely expressed in the honey bee brain, allowing to record neural activity from multiple brain regions. To assess the power of this tool, we focused on the olfactory system, recording simultaneous responses from the antennal lobe, and from the more poorly investigated lateral horn and mushroom body calyces. Neural responses to 16 distinct odorants demonstrate that odorant quality (chemical structure) and quantity are faithfully encoded in the honey bee antennal lobe. In contrast, odor coding in the lateral horn departs from this simple physico-chemical coding, supporting the role of this structure in coding the biological value of odorants. We further demonstrate robust neural responses to several bee pheromone odorants, key drivers of social behavior, in the lateral horn. Combined, these brain recordings represent the first use of a neurogenetic tool for recording large-scale neural activity in a eusocial insect, and will be of utility in assessing the neural underpinnings of olfactory and other sensory modalities and of social behaviors and cognitive abilities.

## INTRODUCTION

Sociality is classified as one of the major transitions in evolution, and animals often form social groups because the benefits of grouping (either direct or indirect) outweigh the costs of breeding independently [1, 2]. The most advanced level of sociality is found in eusocial insect societies [3, 4], and the insect order Hymenoptera (including ants, bees, and wasps) presents the largest number of eusocial species. Among them, honey bees are a classical model for the study of eusocial behavior, as they live in colonies composed of up to 60,000 individuals, consisting of three adult castes (queen, worker, and male). Within the worker caste, honey bees show a clear division of labor with a specialization of roles [5]. The success of honey bee colonies lies in the capacity of all members of the society to behave in a well-organized and context-dependent manner, a social behavior mediated in part by olfactory cues such as pheromones used for communication within the colony [6].

Previous research found that this sophisticated social behavior is associated with higher cognitive abilities in these insects, and in this context, honey bees are a mainstream model for studying higher-order insect cognition such as navigation [7], rule learning [8], social learning [9], dance communication [10], but also olfactory perception and learning [11–13]. However, whether the evolutionary rise of these sophisticated social behaviors is associated with the formation of specific neuronal pathways and structures has not been sufficiently resolved [14, 15]. Progress in understanding these sophisticated behaviors and how social cues are processed and integrated with higher-order cognitive abilities have in part been limited due to the lack of tools allowing to measure neuronal activity simultaneously in different regions of the honey bee brain.

For now, honey bee neuroscience remains limited to the use of conventional neuroanatomy [16], pharmacology [17], electrophysiology [18], and imaging tools [13, 19]. However, the development of neurogenetic tools during the 20^th^ century has provided unprecedented progress in our understanding of the neural basis of behavior in other model species. Among others, this approach allowed to decipher the brain circuits underlying aggressive behavior [20], courtship [21], or memory [22] in a limited set of model species, both vertebrates such as mouse or zebrafish and invertebrates. In insects, the fruit fly *Drosophila melanogaster* has been for many years the leading model for investigating the neural basis of behavior, from gene expression to neural circuits [23–25]. Recently, neurogenetic approaches have been developed in other Diptera such as mosquitoes [26, 27] and in some Lepidoptera [28, 29] to understand specific behaviors such as human-host seeking [30, 31] or insect-plant interactions [32], mainly focusing on their olfactory capabilities. However, genetic methods are particularly difficult to apply in eusocial insects, since genetic transformation rates are low, endogenous promoters for neuronal expressions are unknown and the genetically manipulated, reproductive individuals (the queens) have to be maintained in larger colonies with workers in containments [33].

Despite the difficulty, the recent use of the genome editing tool CRISPR/Cas9 in the honey bee allowed to knock out specific genes, such as the sex-determining *dsx* gene [34], the olfactory co-receptor gene *orco* [35], the gustatory receptor *AmGr3* gene [36], or the *Amyellow-y* gene [37]. Even if these studies evaluated the effect of specific mutations at the neuronal [35] or the behavioral levels [36], versatile genetic tools allowing to investigate the neural basis of a wide range of higher-order social behaviors and learning in honey bees are still missing. The advent of genetically-encoded neural activity sensors, in particular calcium sensors, has represented a major breakthrough in *Drosophila* or mouse research [38–40], but such a critical tool for circuit dissection is still lacking in Hymenoptera.

To this aim, we developed the first pan-neuronal genetic driver in a Hymenopteran model organism, the honey bee, and expressed the calcium indicator GCaMP6f under the control of the honey bee *synapsin* promoter. We characterized its expression pattern in the honey bee brain and evaluated its potential as a functional tool by recording neural activity upon olfactory stimulation. We show that GCaMP6f expression allows to record olfactory responses from multiple brain regions, after simply opening the brain capsule. By using a controlled panel of well-characterized odorants, we show that the recorded signals reveal robust odor coding rules. This new pan-neuronal genetic driver also permits to record neural activity from poorly recorded regions of the brain such as the lateral horn and the mushroom body calyces, and allowed to find that olfactory chemical features are less represented in the lateral horn than in the antennal lobe. This study opens new possibilities for neuroethological research in the honey bee to study the neural basis of advanced social behaviors and cognitive skills.

## RESULTS

This work aimed at expressing the calcium-sensitive protein GCaMP6f in a pan-neuronal manner in the honey bee brain to record neural activity in this social insect.

### Generation of transgenic honey bees

In order to generate a bee with a possibly pan-neuronally expressed calcium sensor, we introduced a synapsin (syn) promoter GCaMP6f expression cassette (*syn-GCaMP6f*) into the genome using the *PiggyBac* transposon system [33]. To generate this *syn-GCaMP6f* expression cassette we cloned the 1 kb promoter region together with the entire 5’ untranslated region (5’ UTR) of the *synapsin* gene from the honey bee (S1 Fig), which we then fused with the coding sequence of the GCaMP6f sensor protein (Fig 1A). We obtained transgenic *syn-GCaMP6f* queens using previously published procedures and a hyperactive transposase [33, 41]. We instrumentally inseminated the queens and reared from only one F0 queen the offspring 2^nd^ generation queens (F1). Only 1% of those F1 queens were carrying the transgene, which we identified from transgene amplifications. The 2^nd^ generation *syn-GCaMP6f* queens produced 50% *syn-GCaMP6f* worker offspring bees, which we used in the following neuroanatomy and imaging experiments.

### Pattern expression of the GCaMP6f in the brain

We first studied the pattern expression of GCaMP6f in the brain using immunostaining against GFP (Fig 1B-D; S2 Fig), the green fluorescent protein which the GCaMP6f sensor contains. We found clear and widespread staining throughout the brain, with strong staining in the antennal lobes (ALs), the optic lobes (OLs), and the mushroom bodies (MBs), suggesting that the GCaMP6f protein is ubiquitously expressed in all major brain structures. In comparison, wild type (WT) bees which do not express the GCaMP6f protein show no staining throughout the brain (S2C Fig).

**Fig 1.**
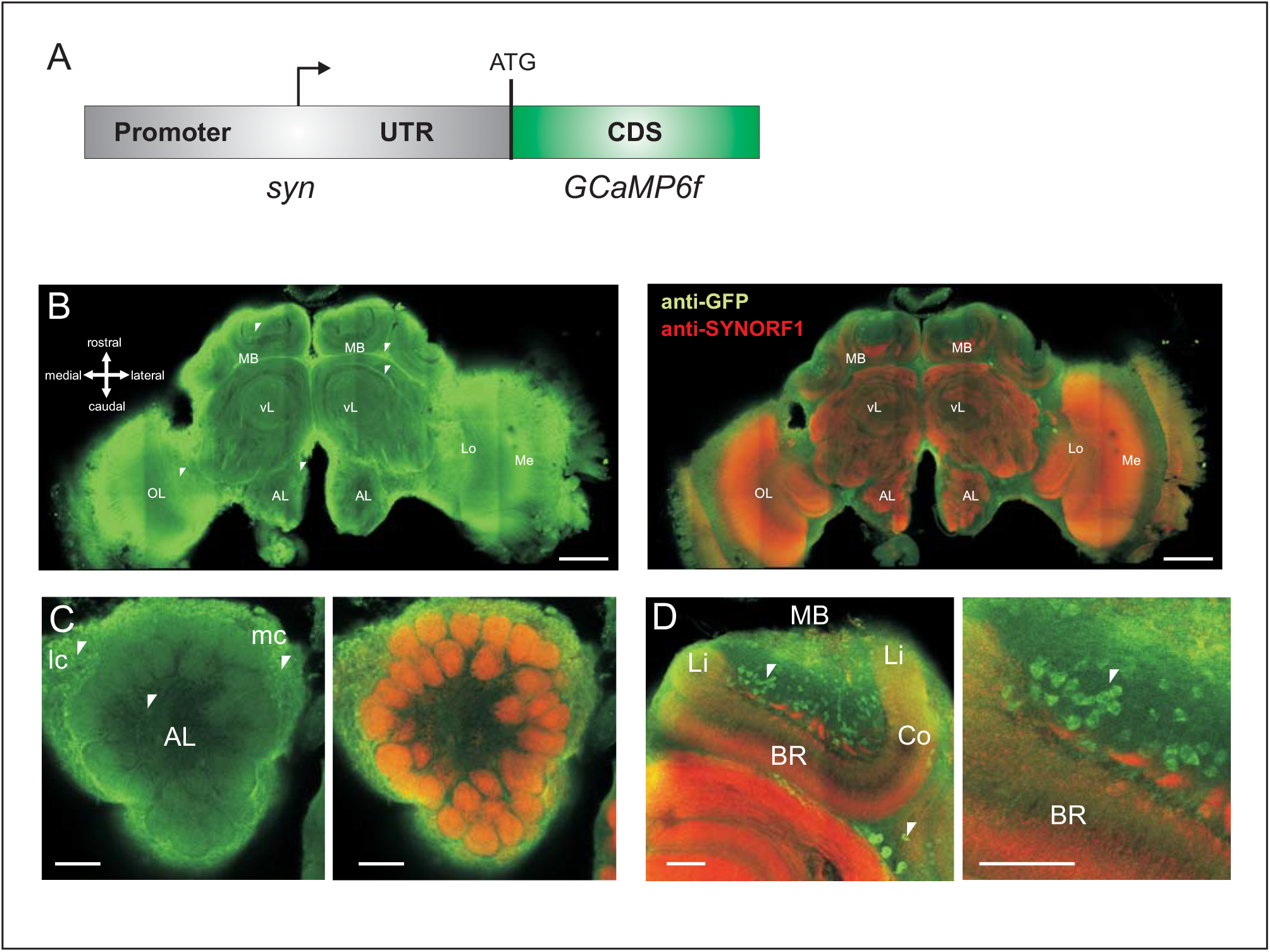
Genetically encoded GCaMP6f and its neural expression. **A-** Scheme of the honey bee *synapsin* promoter *GCaMP6f* expression cassette. CDS: coding sequence; UTR: untranslated region; ATG: translation start. **B-** GCaMP6f expression in the honey bee brain revealed by anti-GFP immunostaining (left, in green). GCaMP6f is widely expressed in the bee brain, including in somata and neural tracts (white arrows). For comparison with *synapsin* expression, an anti-SYNORF1 immunostaining (right, in red) is superimposed on the anti-GFP signal in green. Scale bar = 200 μm **C-** GCaMP6f expression (anti-GFP in green) and *synapsin* expression (anti-SYNORF1 in red) in the antennal lobe. Remarkable and strong expression is observed in the somata of projection neurons and local neurons (near the AL, white arrows). Scale bar = 50 μm. D-GCaMP6f expression (anti-GFP in green) and *synapsin* expression (anti-SYNORF1 in red) in the mushroom bodies with strong expression in some somata of Kenyon cells (in the cup of the calyces, see white arrows). Scale bar = 50 μm. AL: Antennal lobe, MB: Mushroom body, OL: Optic lobe, Lo: Lobulla, Me: Medulla, vL: vertical lobe, lc: lateral cluster of antennal lobe neuron somata, mc: medial cluster of antennal lobe neuron somata, Li: Lip, BR: Basal ring, Co: Collar.

Within neurons of the transgenic bees expressing GCaMP6f, we found evidence of staining in somata (Fig 1B left, S2A-B Fig; white arrows), neuronal processes (see for instance neural tracts around the α-lobe; Fig 1B left, white arrows) as well as dendrites and terminal projections. At the level of the AL (Fig 1C; S2 Fig), lateral and medial clusters of projection neurons and local interneurons somata show strong GCaMP6f immunostaining. In addition, glomeruli seem homogeneously stained: staining in the cortex suggests expression in olfactory sensory neurons (OSNs) terminal projections, whereas staining in the glomerulus core suggests expression in projection neurons (PNs) dendrites and possibly local neurons. Finally, the clearest expression of GCaMP6f was found in somata at the level of the calyces of the MB (Fig 1D; S2B Fig). In these structures, different neuron types are stained, with somata of class I Kenyon Cells in the cup-shaped calyces and class II Kenyon Cell somata lying outside the calyces (see white arrow in Fig 1D; S2B Fig) [42, 43]. These staining patterns were similar in the different bees we used for the immunostaining (S2A-B Fig). We found very broad expression in some groups of neurons like the somata of antennal lobe neurons (Example shown in S3A Fig for a somata cluster of local/projection neurons in caudo-lateral position) and at the same time heterogeneous staining at the level of the Kenyon cells within MB calyces. We also found expression of GCaMP6f in neurons outside of the central brain, for example in the photoreceptors of the ocelli (S3B Fig).

We next asked whether the cloned synapsin promoter used for GCaMP6f expression drives the same expression pattern as the one of the *synapsin* gene. To answer this question, we performed a complementary immunostaining targeting the synapsin protein (Fig 1B right, anti-SYNORF1 in red). We found co-localized staining of GCaMP6f and synapsin within AL glomeruli both in the cortex and the core (Fig 1C right), demonstrating co-expression within AL neurons. This experiment also shows differences between GCaMP6f and synapsin stainings. The synapsin immunostaining allows to clearly visualize presynaptic zones, for instance in AL glomeruli or microglomeruli in the MB calyces, but is not present in the somata unlike GCaMP6f (Fig 1D, white arrows). This result indicates that even if GCaMP6f is expressed in cells expressing synapsin (in theory all neurons), these 2 proteins appear to be partially differently localized within neurons.

We conclude that GCaMP6f was widely expressed in the honey bee brain, one of the prerequisites for whole-brain functional recordings of neural activity.

### Calcium imaging

#### Neural recordings in the whole brain

GCaMP6f expression in neurons was then used to record neural activity in the honey bee brain (Fig 2). As a proof of concept for the use of these bees in neuroethological research, we decided to focus here on olfactory information processing, as olfaction is the best-understood sensory modality in honey bees. Right after placing the bee in a recording chamber and opening its head capsule, we first presented a few standard odorants while recording the whole brain surface accessible under the microscope objective. Odorant presentations triggered a clear GCaMP fluorescence change from several brain regions (Fig 2A), including the primary olfactory centers (the antennal lobes, ALs) and both higher-order centers (the lateral horns, LHs and the mushroom bodies, MBs), known for their role in olfactory processing. The use of GCaMP-expressing bees allowed us to record neural activity simultaneously in all these structures (Fig 2B), while the presentation of an unscented air control did not induce strong activity. Response amplitudes were stronger in the AL (Fig 2C left) than in the LH (Fig 2C middle) or in the MB (Fig 2C right). In all regions, the recorded calcium signals showed a biphasic response, with a fluorescence increase upon odor presentation followed by a long undershoot. Such biphasic signals are reminiscent of calcium signals previously recorded using bath-applied calcium-sensitive dyes [44, 45]. This experiment represents the first recording of neural activity using neurogenetic tools outside of Diptera.

**Fig 2.**
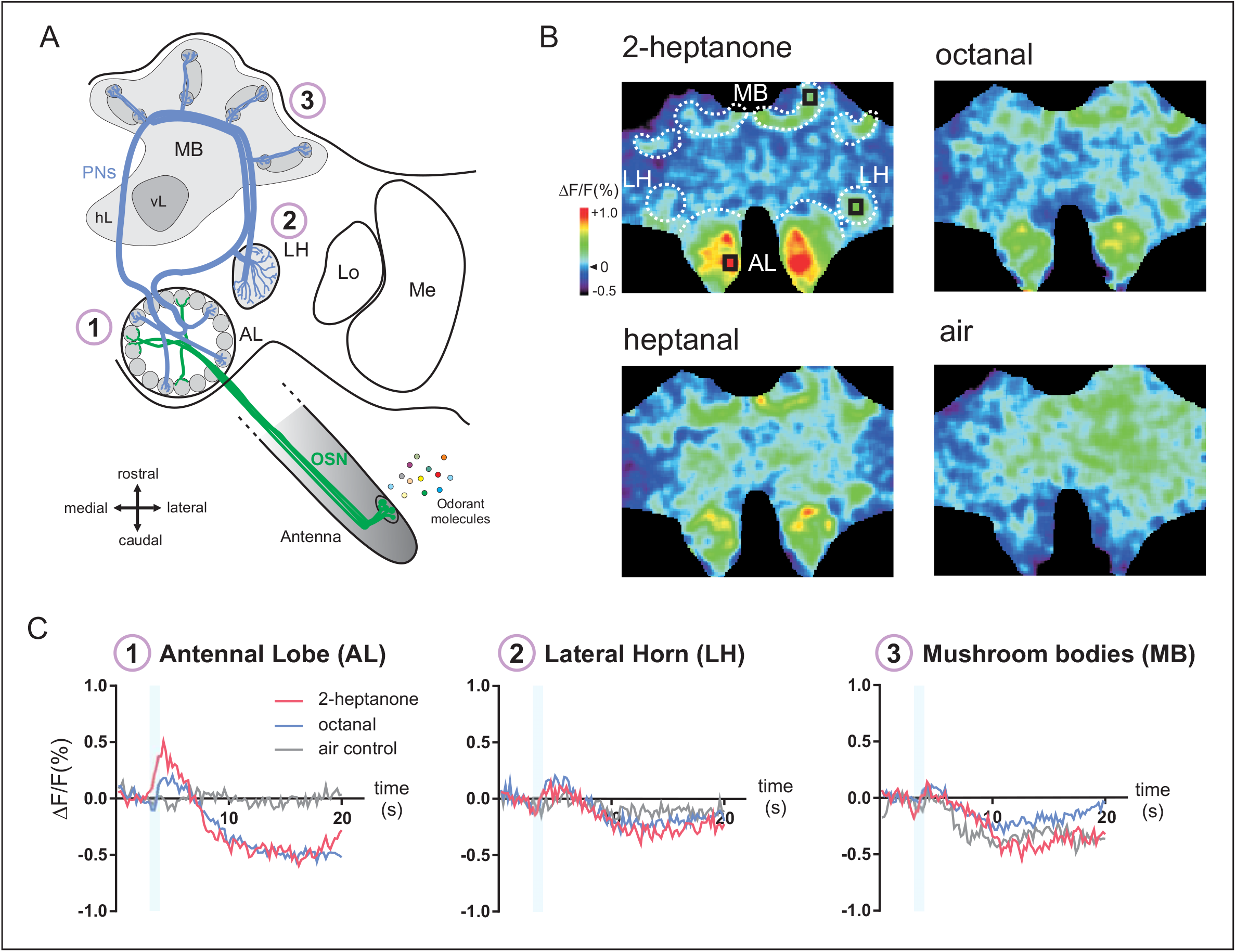
Neural activity in the whole brain of the honey bee. **A-** Hymenopteran olfactory pathway (adapted from [45]). Odorant molecules are detected by olfactory sensory neurons (OSNs) on the antenna, which send olfactory information to the antennal lobe (AL). Then, projection neurons (PNs) convey information to higher-order centers, the mushroom bodies (MB) and the lateral horn (LH). Lo: Lobula, Me: Medulla, vL: vertical lobe, hL: horizontal lobe. **B-** Calcium signals in the whole brain evoked by 3 different odorants (2-heptanone, octanal, heptanal) and the air control. Relative fluorescence changes (Δ*F*/*F* [%]) are presented in a false-color code, from dark blue (minimal response) to red (maximal response). **C-** Time course of odor-evoked responses (Δ*F*/*F* [%], taken from the black squares shown in B) for one individual to the presentation of 2-heptanone (in blue), octanal (in pink), and the air control (in grey), simultaneously recorded in the AL (left), the LH (middle) and the MB (right). The calcium signals show a biphasic time course, with a fluorescence increase upon odorant presentation (blue bars) followed by a long-lasting fluorescence undershoot, in all brain regions.

#### Neural recordings in the antennal lobe (AL)

We then asked if the signals recorded using this genetically-encoded calcium sensor represent neural signals that are meaningful in terms of sensory coding, focusing first on the primary olfactory center, the antennal lobe (AL), since olfactory coding was best studied in this structure. To this aim, we presented 16 aliphatic odorants used in previous imaging studies either after bath-application of a cell-permeant calcium-sensitive dye (Calcium-green 2-AM, OSN recordings) [45] or after insertion of a migrating calcium-sensitive dye within a neural tract (Fura-2 dextran, PN recordings) [19]. The response patterns to the 16 aliphatic odors differed systematically according to their functional group (primary or secondary alcohols, aldehydes, or ketones) or their carbon chain length (from 6 to 9 carbons) (Fig 3A). As observed with other calcium reporters, the presentation of each odorant induced a biphasic signal in a different set of AL glomeruli while presentation of the air control or no stimulation did not induce such response (Fig 3B, S4A Fig; S5 Data; S9 Data). Moreover, WT bees did not show any signal in response to odor presentation (S4B Fig). The averaged AL response amplitudes were significantly different from the air control for all 16 odors (Fig 3C; S1 Data; *n* = 11 honey bees, RM-ANOVA, odor effect *F*_16,160_ = 12.49, *p* = 8e-21, post hoc Dunnett tests *p* < 0.0097). Signal amplitudes differed according to the odor (odor effect, *p* = 8e-21), and were related to the quantity of molecules in headspace, since odor-evoked responses were highly correlated with the vapor pressure of the odorants (Fig 3D, *n* = 11 honey bees, *R*^2^ = 0.84, Fisher test *F*_1,14_ = 73.72, *p* = 6e-7). Such a correlation was also found in previous studies recording at AL input (OSNs) [45] or output (PNs) [19]. This strong dependence explains why odor-evoked intensities recorded using genetically encoded GCaMP6f in this study are highly correlated with odor-evoked intensities recorded in OSNs (S5A Fig, *R*^2^ = 0.84, *F*_1,14_ = 74.23, *p* = 6e-7) or in PNs (S5B Fig, *R*^2^ = 0.78, *F*_1,14_ = 49.73, *p* = 6e-6).

**Fig 3.**
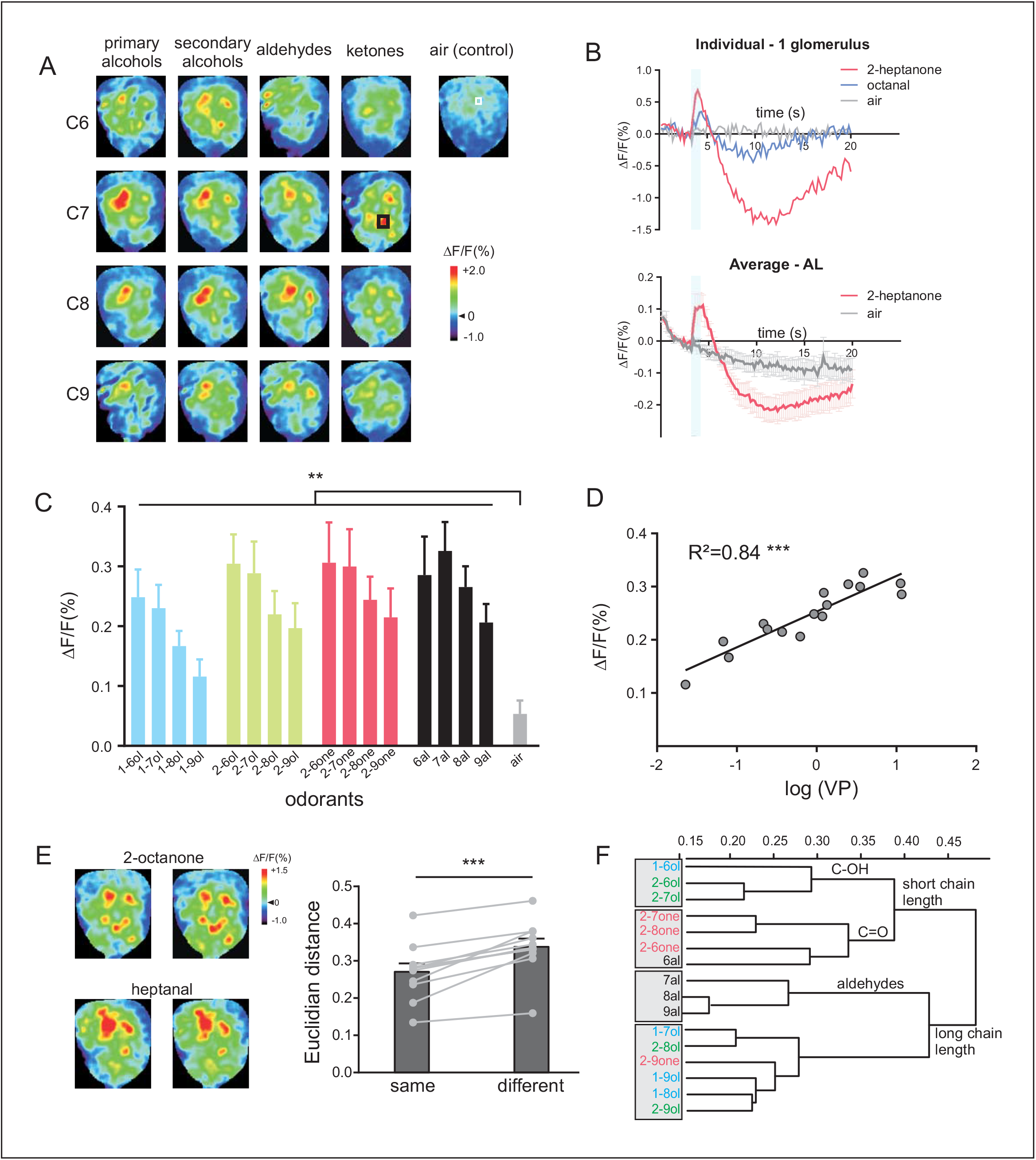
Neural activity recorded in the antennal lobe (AL) **A-** GCaMP6f calcium signals in the antennal lobe evoked by a panel of 16 odorants varying systematically according to their carbon chain length (C6–C9) and their functional group (primary and secondary alcohols, aldehydes, and ketones). Different odorants induce different glomerular activity patterns. The map shows the whole amplitude of the response including both positive and negative components (see text for calculation). **B-** Example time courses (top, taken from the black square shown in A in C7 ketones) and average time courses (bottom, *n* = 11 honey bees) of odor-evoked responses (Δ*F*/*F* [%]) recorded in the AL to 2-heptanone (in red), octanal (in blue), and to the air control (in grey). **C-** Amplitude of calcium responses (Δ*F*/*F* [%]) to the 16 aliphatic odorants and to the air control. All odorants induce significant activity in comparison to the air control (*n* = 11 honey bees, ** *p* < 0.0097). The intensity of the odor-induced response was obtained by averaging three consecutive frames at the end of the odor presentation (frames 19-21) and subtracting the average of 3 frames during the second, negative component of the signal (frames 49-51). **D-** Amplitude of calcium responses (Δ*F*/*F* [%]) as a function of odorant vapor pressure (in log units). The linear regression shows a significant correlation (*R*^2^ = 0.84, *** *p* = 6e-7). **E-** Different presentations of the same odorants (2-octanone and heptanal) show similar glomerular patterns in the AL (left). Dissimilarity measures (Euclidian distance, right) between representations of the same or of different odorants. Activity maps are more similar (shorter distances) when the same odorant is presented (*** *p* = 1.1e-4), showing, as expected, a clear odor coding in this structure. **F-** Cluster analysis showing similarity relationships among odorants (Ward’s classification method). Functional groups are shown in different colors: primary alcohols in blue, secondary alcohols in green, aldehydes in black, and ketones in red. The analysis shows a first separation between odorants with short and long carbon chain lengths. Odorants with a short carbon chain are then subdivided into alcohols (primary and secondary, C-OH functional group) and ketones/aldehydes (C=O functional group). The data underlying the graphs shown in the figure can be found in S1 Data and S5 Data.

We then evaluated odor coding in the GCaMP bees by comparing similarity relationships among odorants, by calculating Euclidian distances between response maps obtained for the different odor pairs (S1 Data). First, we confirmed that within each bee, each odorant evokes a specific activity pattern, as distances were lower for different presentations of the same odorant than for the presentation of different odorants (Fig 3E, *n* = 11 honey bees, paired *t*-test *t* = 6.15, *p* = 1.08e-4). This result fits with previous calcium imaging data both at AL input and output [46, 47]. We then performed a hierarchical cluster analysis using Euclidian distances between odorants. We found a first segregation between odorants with long chain lengths and odorants with short chain lengths (Fig 3F, C6-C7 vs. C8-C9). Within the shorter chain length group, a further separation between functional groups appeared based on the oxygen moiety (C=O for aldehydes and ketones vs. C-OH for alcohols). Within the longer chain length group, aldehydes were separated from the other odorants. This analysis shows a clear coding depending on the chemical structure of the odorants, and with remarkably similar results as in previous studies using non-transgenic approaches [19, 45]. Accordingly, inter-odorant distances found in the GCaMP bees were highly significantly correlated with those measured previously in OSNs using calcium dye bath application (*R*^2^ = 0.53, Mantel test *p* = 1.0e-4) or measured in PNs using dye injection (*R*^2^ = 0.44, Mantel test *p* = 1.0e-4). The correlation coefficients were not significantly different (*R*^2^ = 0.53 vs. *R*^2^ = 0.44, Fisher test, *z* = 0.98, *p* = 0.33, NS), suggesting the possibility that both OSN and PN contribute to the signals recorded in the AL using genetically encoded GCaMP6f.

Finally, we asked if odorant response maps recorded with the GCaMP bees relate to bees’ actual perception of the odorants. We thus correlated the Euclidian distances measured between all odor pairs in this study with distances measured in an appetitive conditioning experiment (S8 Data) [48]. We found that behavioral distances can be predicted using signals recorded with genetically encoded GCaMP6f (S6 Fig; *R*^2^ = 0.34, Mantel test *p* = 1.0e-4). Thus, odorants evoking similar activity patterns in the AL are treated as similar by honey bees in their behavior, a further confirmation that GCaMP6f expression allows to record meaningful neural signals.

#### Neural recordings in the lateral horn (LH)

We then studied olfactory information processing in higher-order brain centers, first focusing on the LH. All the aliphatic odorants used in this panel induced calcium signals in the LH (Fig 4A). Odor-evoked signals systematically followed a biphasic time course, like that recorded in the AL (Fig 4B; S7 Fig; S6 Data). In the LH, all but one of the presented odorants induced a response that was significantly higher than to the air control (S2 Data, *n* = 8 honey bees, RM-ANOVA, odor effect *F*_16,112_ = 8.091, *p* = 0.0014, post hoc Dunnett tests, *p* < 0.04 except *p* = 0.09 for 2-hexanol). As in the AL, response amplitudes correlated with odorants’ vapor pressures (*R*^2^ = 0.49, *F*_1,14_ = 13.87, *p* = 0.0023). However, this correlation appeared to be weaker in the LH with a near-significant difference between the two correlation coefficients (*R*^2^ = 0.84 in the AL vs. *R*^2^ = 0.49 in the LH, Fisher test, *z* = 1.76, *p* = 0.078). In other words, response intensity in the LH was not a simple product of the number of volatile molecules in the odor puff, possibly revealing a yet undocumented gain control mechanism in the LH.

**Fig 4.**
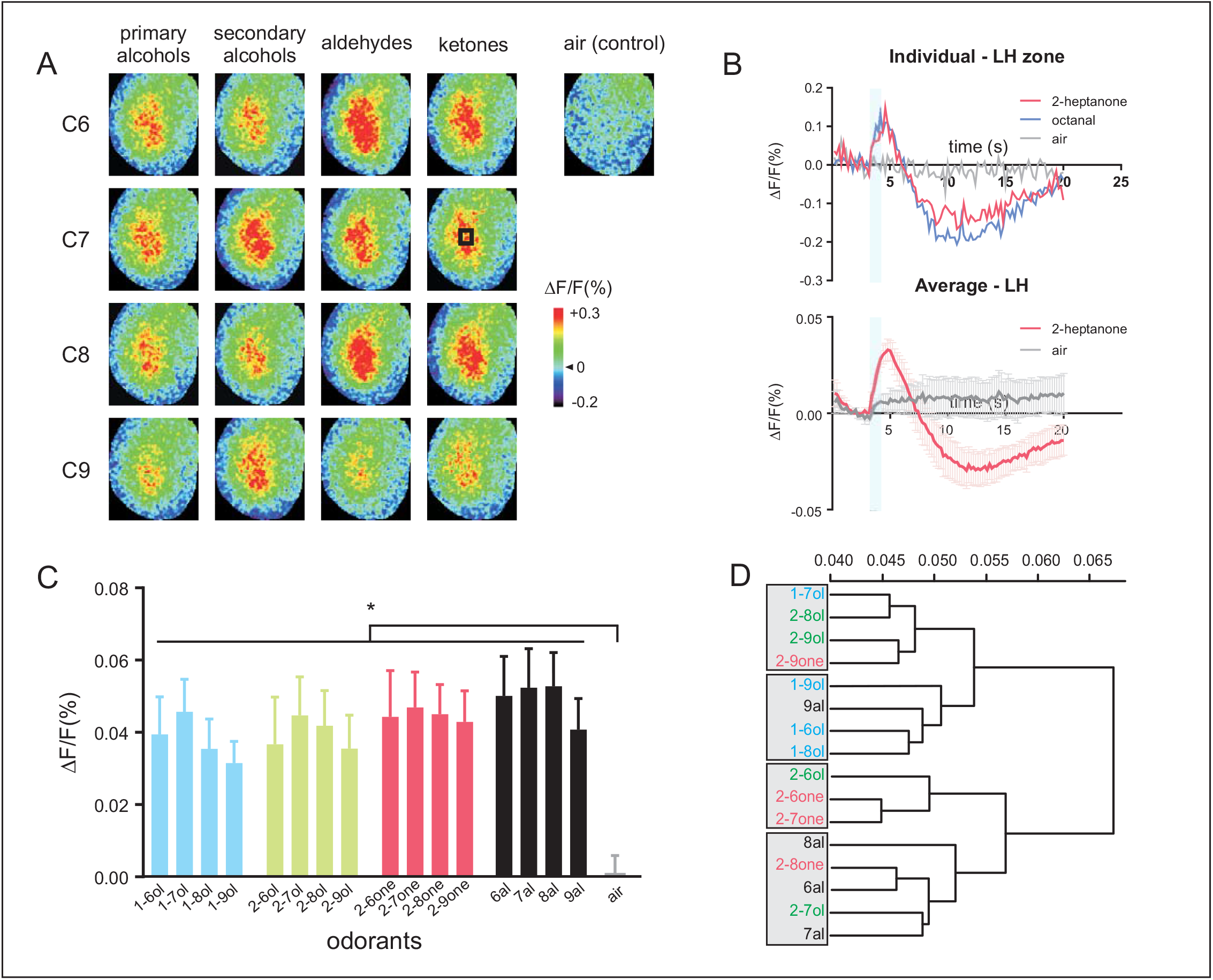
Neural activity recorded in the lateral horn (LH) **A-** Calcium signals in the lateral horn evoked by the same panel of 16 odorants. Different odorants induce different activity patterns in the LH. **B-** Example time courses (top, taken from the black square shown in A in C7 ketones) and average time courses (bottom, *n* = 8 honey bees) of odor-evoked responses (Δ*F*/*F* [%]) recorded in the LH (in the black square shown in A) to 2-heptanone (in red), octanal (in blue), and to the air control (in grey). The calcium signals also show a biphasic response, with a fluorescence increase upon odor presentation (blue bar) followed by a long undershoot. **C-** Amplitude of calcium responses (Δ*F*/*F* [%]) to the 16 aliphatic odorants and to the air control. All odorants induce a significant activity in comparison to the air control *(n* = 8 honey bees, * *p* < 0.04). **D-** Cluster analysis showing similarity relationships among odorants (Ward’s classification method). The data underlying the graphs shown in the figure can be found in S2 Data and S6 Data.

We then compared similarity relationships (Euclidian distances among odorants, S2 Data) measured in the LH and in the AL and found a weak but still significant correlation (*R*^2^ = 0.07, Mantel test *p* = 0.023). This suggests a transformation of odor similarity relationships between the two structures. A cluster analysis performed on Euclidian distances in the LH confirmed this finding. The analysis roughly segregated alcohols (-OH moiety) on the one side and aldehydes/ketones (=O moiety) on the other. However, different rules apply here, as the two longest chain molecules nonanal and 2-nonanone clustered with the former group, while the shortest chain secondary alcohols, 2-hexanol and 2 heptanol clustered with the latter group. Thus, the study of odor coding in the LH using GCaMP bees suggests a less clear dependence on chemical features than in the AL.

Nevertheless, Euclidian distances measured in the LH correlated significantly with behavioral distances measures [48] (*R*^2^ = 0.12, Mantel test *p* = 0.0021), demonstrating that odorants evoking similar activity patterns in the LH are treated as similar by honey bees in their behavior.

#### Neural recordings in the mushroom bodies (MB)

Lastly, we turned to the second higher-order center, the mushroom bodies (Fig 5, S3 Data), a multisensory integration center in the honey bee brain. Calcium signals upon odorant presentation were observed at the level of the calyx lip. Calcium signals systematically showed a biphasic time course (Fig 5B; S8 Fig; S7 Data) like that recorded in the AL and in the LH, but in all cases with a small first component and a much stronger second component. In this structure, the five presented odorants induced significant activity in the MB calyx in comparison to the air control (Fig 5C; *n* = 6 honey bees, RM-ANOVA, odor effect *F*_5,25_ = 7.99, *p* = 0.0033, post hoc Dunnett tests *p* < 0.05, except 2-heptanone with *p* = 0.07). However, the signal-to-noise ratio was considered too low for further analysis of inter-odor relationships.

**Fig 5.**
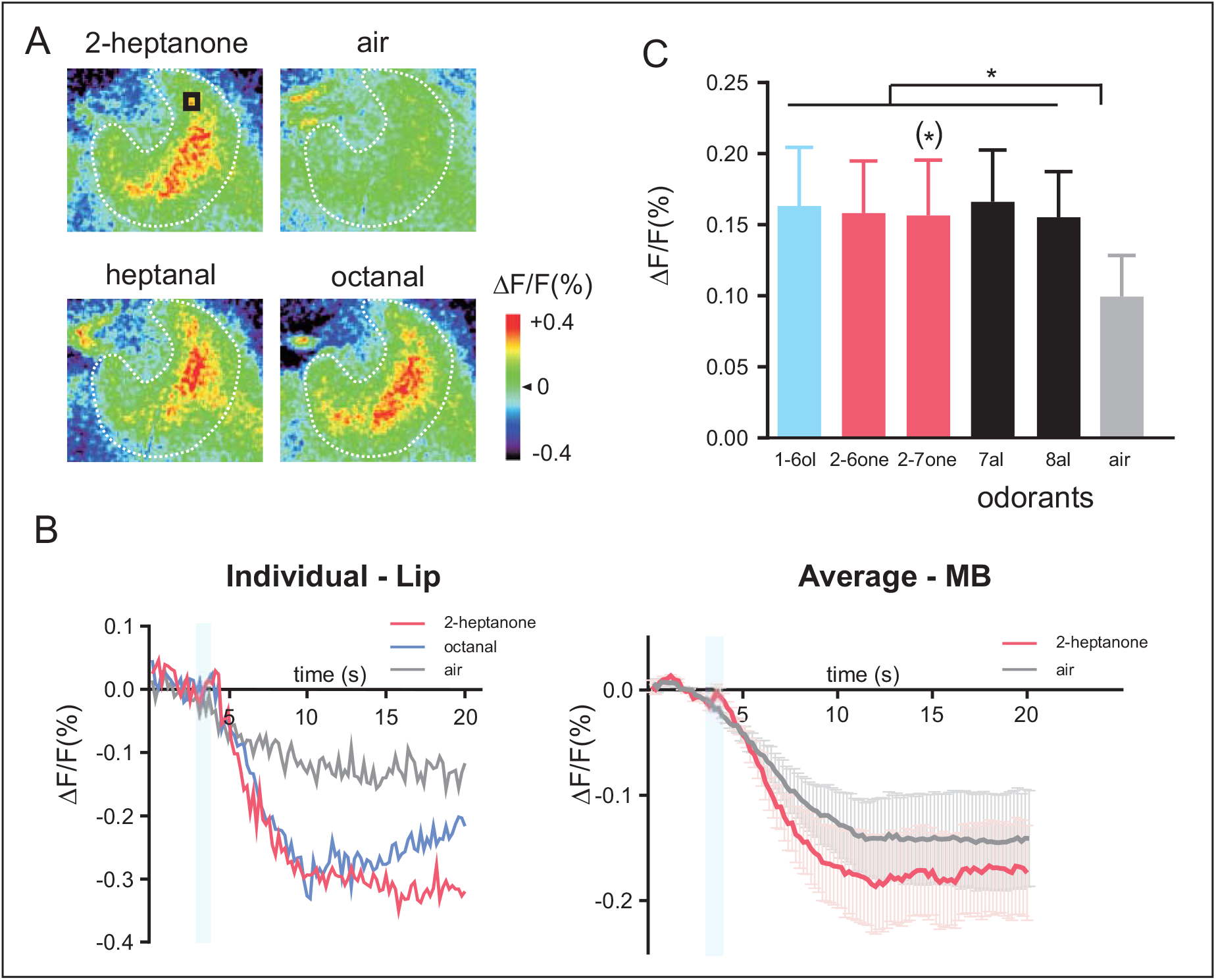
Neural activity recorded in the mushroom bodies (MB) **A** - Calcium signals in the calyces of the mushroom bodies in response to the presentations of 2-heptanone, heptanal, octanal, and the air control. **B-** Example time courses (left, taken from the black square shown in A in 2-heptanone) or average time courses (right, *n* = 6 honey bees) of odor-evoked responses (Δ*F*/*F* [%]) recorded in the lip of the MB (black square shown in A) to 2-heptanone (in red), octanal (in blue), and to the air control (in grey). Calcium signals are weaker here than in other structures, with a smaller first component and a larger second component. **C-** Amplitude of calcium responses (Δ*F*/*F* [%]) to the 5 tested odorants and to the air control. All odorants induce significant activity in comparison to the air control (*n* = 6 honey bees, * *p* < 0.05, except 2-heptanone with *p* =0.07). The data underlying the graphs shown in the figure can be found in S3 Data and S7 Data.

#### Neural recordings of responses to social pheromones

A condition for using GCaMP bees for unraveling the neural basis of social communication is the ability to record robust responses to pheromonal odorants. We focused again on the LH and presented a panel of honey bee pheromonal odorants, including both volatile (IPA, from the alarm pheromone and ocimene, a volatile brood pheromone) and non-volatile compounds (queen and brood pheromone compounds). Calcium signals were visible for all compounds in the LH (S4 Data, Fig 6A), although the response intensity was significantly higher from the air control (Fig 6B, *n* = 7 honey bees) for volatile compounds (*t*-tests, *t* > 4.35, *p* < 0.005) and for queen pheromone compounds (Friedman ANOVA, odor effect *F*_11_ = 30.23, *p* = 0.0008; *t*-tests or Wilcoxon tests, *t* > 2.48, *p* < 0.05 except for 9-ODA with *t* = 1.8 and *p* = 0.11) but not for brood pheromone compounds (*t*-tests *t* < 2.09, *p* = NS).

**Fig 6.**
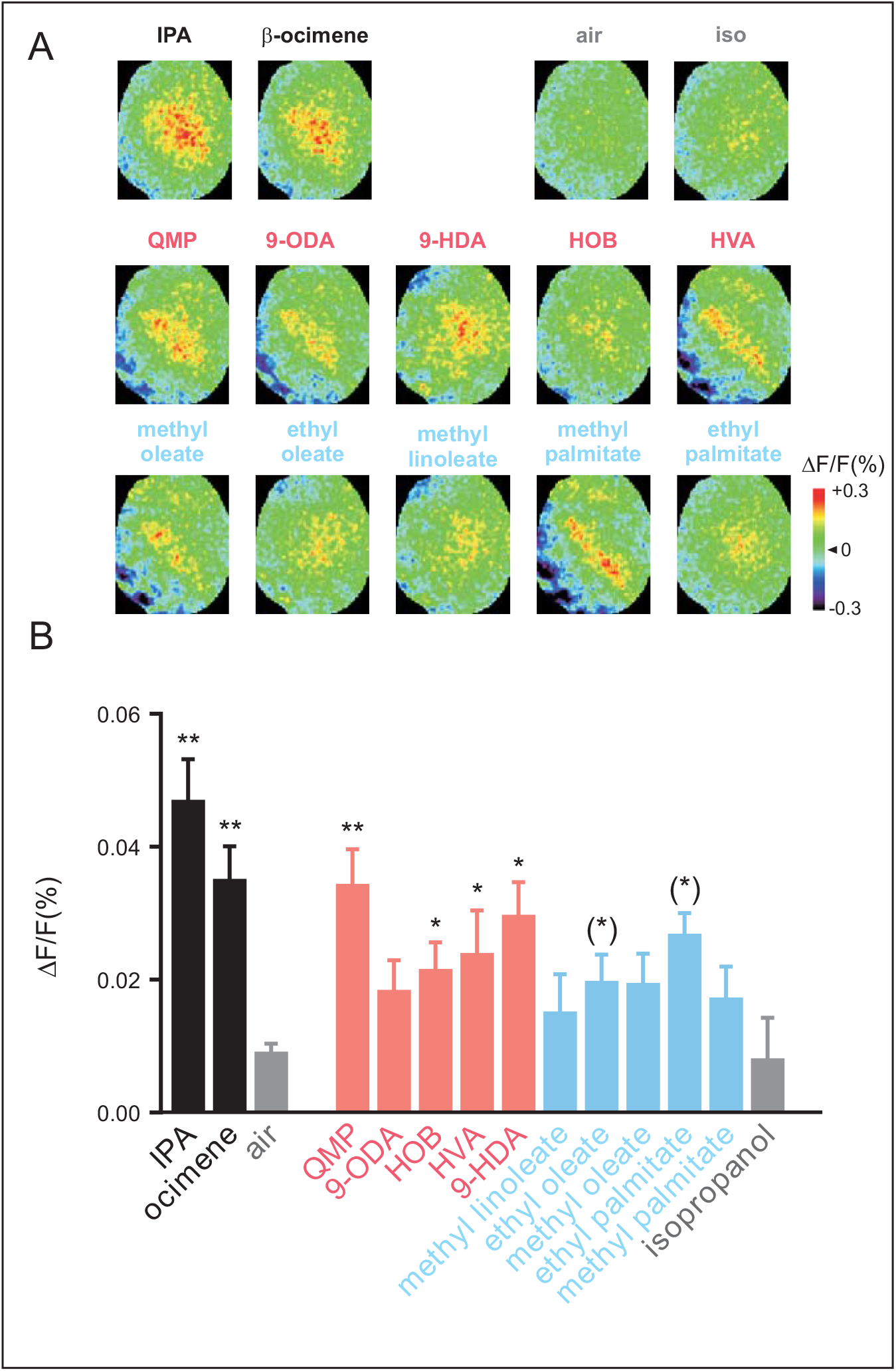
Neural responses to pheromonal odorants recorded in the lateral horn. **A-** GCaMP6f calcium signals in the lateral horn evoked by a panel of pheromonal compounds, both volatile (IPA and ocimene in black) or not volatile (queen pheromone compounds in red and brood pheromone compounds in blue) and to the controls (isopropanol and air). Different odorants induce activity patterns. Relative fluorescence changes (Δ*F*/*F* [%]) are presented in a false-color code (see colorbar), from dark blue (minimal response) to red (maximal response). **B-** Amplitude of calcium responses (Δ*F*/*F* [%]) to the 12 tested odorants and to the controls (air and isopropanol). Volatile odorants and queen pheromone compounds induce significant activity in comparison to the controls (*n* = 7 honey bees, * *p* < 0.05). The data underlying the graphs shown in the figure can be found in S4 Data.

## DISCUSSION

We developed a line of honey bees with pan-neuronal expression of a genetically-encoded calcium sensor allowing to record neural activity from honey bee neurons. To our knowledge, this represents the first neurogenetic tool for recording neural activity in a non-dipteran insect and eusocial insect. Neurogenetic tools, especially the use of genetically-encoded calcium indicators (GECIs), have been widely developed in both vertebrates [40, 49] and invertebrates [50] and supported major progress in our understanding of the neural basis of behavior. In insects, such possibility was limited until now to a few Dipteran species (fruit flies and mosquitoes [26]) and was lacking in other insects with different lifestyles. The present development in a social Hymenoptera represents a major progress for the neuroethology of social behavior but also a significant achievement in a species like the honey bee in which neurogenetic tools are difficult to establish due to its peculiar reproductive biology and the lack of knowledge of appropriate promoter sequences driving the expressions.

To develop our pan-neuronal calcium indicator, we expressed GCaMP6f under the control of the promoter of the gene coding for synapsin, a vesicle–associated presynaptic protein found throughout the honey bee brain [51]. This strategy was successful, as honey bees displayed broad labeling with GCaMP6f in the major brain structures of the nervous system at sufficient amounts for performing calcium imaging. Our neuroanatomical data confirm significant expression throughout the bee brain (optic lobes, antennal lobe, mushroom bodies, and all other regions of the protocerebrum), but we noticed substantial cell-to-cell variation in some regions, for instance in the cup of the mushroom body calyx, which contains the somata of type I Kenyon cells (see Fig 1C): there, a strong contrast appeared among neurons, suggesting differences in the level of GCaMP expression among Kenyon cells or even some Kenyon cells lacking GCaMP expression. The GCaMP staining pattern did not correspond to any of the Kenyon cell subtypes previously described based on their differing gene expression profiles [52]. In contrast to the synapsin protein, GCaMP6f is located in the entire neuron, since we detected staining in the somata (Fig 1C), the neurites, and the synaptic regions. This widespread expression could represent a great advantage for performing functional recordings in the bee brain.

To assess the power of this new tool for studying brain activity, we focused on the olfactory system, which is the best-known sensory system in this animal. We compared the results obtained using the GCaMP6f bees with previous results acquired using conventional dye injections or bath application [19, 45]. For the first time, we could observe the simultaneous activation of the three main brain structures involved in insect olfaction, the antennal lobe, the lateral horn, and the mushroom body calyx.

While odor stimuli elicited responses well confined to these structures, also pure air stimuli produced neuronal activation but distributed over further central brain regions. These could be responses of various mechanosensory neurons, which in principle should be largely suppressed by the olfactometer keeping a constant airflow. However, small fluctuations during the switching phases are inevitable. Besides neuropils that process mechanosensory information from the Johnston’s organ, such as the dorsal lobe, the posterior protocerebral lobe, and the subesophageal zone [53], also MB [54] and LH [55] are known to integrate mechanosensory information and recently mechanosensitivity has also been reported in olfactory neurons [56, 57].

The possibility to record from different brain structures at the same time is a first and important step towards the long-term goal to understand brain circuits responsible for bees’ sophisticated social behaviors and cognitive abilities, that may unravel yet undescribed sensory and/or behavior-related pathways.

To follow this, we first validated our tool using individual structures and we confirmed its validity for measuring relevant neural activity with regard to bees’ behavior. Recordings obtained using the GCaMP6f-expressing honey bees revealed the same olfactory coding rules in the AL as in previous studies [19, 58]. Thus, odor-induced activity followed the same intensity rule (depending on the odorant vapor pressure – Fig 3D), and inter-odorant neural distance measures were organized according to odorants’ chemical features, chain length, and functional group. Accordingly, calcium imaging measured with the GCaMP-expressing bees allowed to predict how similarly honey bees perceive these odorants, since inter-odor relationships among GCaMP response maps correlated with inter-odor behavioral distances previously measured in an associative conditioning experiment [48]. This result, even if limited to a single sensory modality, is a strong confirmation of the biological pertinence of the signals measured with this new neurogenetic tool.

In order to demonstrate that the activity can be followed to higher-order brain structures, we studied the role of the lateral horn in encoding the chemical features of our odorants. The lateral horn is a poorly studied structure in Hymenoptera, which could only be recorded previously using invasive dye injection [59]. Importantly, while previous recordings only measured from a limited subpopulation of projection neurons in this structure (the l-ALT), the GCaMP6f-expressing honey bees allowed recording from the whole structure. The recordings in response to the 16 aliphatic odorants showed that odor coding in this higher-order center is less clearly influenced by odorant quantity (*i.e*. lower response amplitude differences among odorants) or by the chemical structure of the odorants than in the AL. These first results suggest the existence of an additional gain control mechanism in this structure, fitting with current discussions on its more diverse role in *Drosophila* [60]. This includes a possible coding of hedonic odorant valence rather than chemical features [55, 61], a segregation of food odors from pheromones [62], a central involvement in courtship behavior [63], or a multisensory integration of olfactory, visual, mechanosensory, and gustatory information [55, 61]. The tool presented here will allow for comparable studies also in the honey bee.

Calcium signals were also recorded in the mushroom body calyx, and odorants clearly activated the lip, the olfactory input region of the calyx, as observed previously with a bath-applied dye [64]. The signal amplitude was generally low, which reproduces the known sparseness of the MB odor code [65]. Together with the random arrangement of PN terminals and KC dendritic arbors in the lip, this prevented the observation of odor-specific response maps at this point. However, advanced 3D microscopy techniques [66] will benefit from this neurogenetic marker to unravel coding mechanisms also in this neuropil, as experiments in *Drosophila* promise [67].

This new tool will provide a range of advantages over previous solutions. We expect that, thanks to the genetically-encoded calcium sensor, it will facilitate imaging from other members of the colony, like queens and drones, in contrast to what has been done until now (an exception is [68]). Furthermore, it will also allow to extend the study of sensory processing to other sensory modalities than the most commonly studied olfaction and vision. For instance, the study of mechanosensory information processing notably used during the waggle dance remains limited [57, 69] and should be facilitated using GCaMP-expressing honey bees. Lastly, transgenic bees could be instrumental for studying neural signals in response to multisensory inputs, for instance in the mushroom bodies, the higher-order center known for its role in multimodal sensory integration [70].

The future of insect neuroethology beyond the use of model organisms like *Drosophila* will depend on the further development of neurogenetic tools in species like honey bees, with their rich behavioral repertoire involving intricate social behaviors as well as elaborated cognitive skills. First, the use of the CRISPR-Cas9 strategy, already developed in the honey bee [34], possibly allows target specific insertion with an integration rate higher than the rate achieved with the PiggyBac transposon system. It should be noted that once a transgenic queen is produced, the line can be kept for longer periods of time by regularly raising new queens from the eggs laid by the initial queen. This procedure involves relatively standard beekeeping practices. The future neurogenetic tools will also have to improve the amplitudes of the calcium responses, with a better signal-to-noise ratio, a necessary condition to study sensory coding in higher-order centers. A potential strategy would be to amplify the GCaMP expression using the UAS-Gal4 or the Q binary expression systems commonly used in fruitflies and mosquitoes [71]. Lastly, a better resolution will have to be achieved using cell-type-specific gene expression to visualize (*i.e*. using fluorescent proteins such as GFP) or monitor activity (using GECIs as here) from specific neuronal populations. The same precision shall then be exploited to silence or activate specific neuron populations, allowing a temporal control of neural function. We hope that the new line of GCaMP-expressing bees will represent the first step in this direction.

## MATERIAL AND METHODS

### *PiggyBac* transgenesis

The upstream promoter region together with the three exons of the 5’ UTR of the honey bee *synapsin* (*syn*) gene (LOC551737; assembly Amel_4.5, gene annotation 104) were successively cloned (S1 Fig). This single fragment (1811 bp) was inserted into the PBac plasmid [33] using the *Asc*I/*Nco*I restriction sites. The coding sequence of *GCaMP6f* was inserted downstream of the promoter region using *Nco*I/*Mss*I restriction sites that finally resulted in the PBac [*syn-GCaMP6F*]Am plasmid. Requests for the PBac *[syn-GCaMP6F]Am* plasmid should be directed to and will be fulfilled by Martin Beye. Egg injections, larvae and queen rearing were performed following standard methodology by employing the hyperactive transposase which we have customized for the honey bee [33, 41]. A total of 4231 honey bee eggs not older than 1.5 h were injected, with 82% survival after 24h. Following the hatching, 34 eggs were reared to queens and only 7 out of the 34 were expressing the GCaMP6f transgene. Semen from wild type drones was used for instrumental insemination of queens. Inseminated queens were laying fertilized, usually female-determined eggs that were reared either to queens or worker bees. Transgenic queens were identified by amplification of the transgene in the offspring [41] using oligonucleotide primers (CCACACCTCCCCCTGAACCTGAAAC and GAGGTAAGAATAAACATTGTTGGTC) targeting the PBac sequence. Only one inseminated queen was kept for this study, and 2^nd^ generation queens were reared from its offspring. Only 1% of the 2^nd^ generation queens were carrying the transgene suggesting that only a fraction of germ cells from the 1^st^ queen carried the transgene. However, 50% of the offspring of the transgenic 2^nd^ generation queens were carrying the transgene, and workers were used for the anatomical staining and the calcium imaging recordings.

### Anatomical immunostaining

To visualize the expression pattern of the GCaMP6f in the honey bee brain, transgenic workers were caught at emergence and were maintained in an incubator in the dark at 34°C for 2 weeks (*i.e*. the same age as for calcium imaging). The brains of workers were dissected and immediately immersed in cold 1% zinc formaldehyde in PBS (ZnFa 1% [72]) and kept overnight at 4°C. Brains were then washed six times in PBS (10 min each), permeabilized in PBS containing 1% Triton X-100 for 30 min, and pre-incubated 3h in PBS containing 0.3% Triton and 1% BSA (Bovine Serum Albumine in PBS, #37525, ThermoScientific). To stain *GCaMP6f* and *synapsin* (as background staining), the brains were then incubated in PBS containing 0.3% Triton and 0.1% BSA with rabbit polyclonal anti-GFP (Thermofisher, #A-11122, France) at 1:100 dilution and mouse monoclonal anti-SYNORF1 (DHSB, #3C11, US) at 1:100 dilution for 7 days. Brains were then washed 6 times in PBS containing 0.3% Triton and incubated in secondary antibodies directed against rabbit coupled to Alexa 488 (Thermofisher, #A11122, France) and against mouse coupled to Alexa 555 (Thermofisher, #A-21147, France) both diluted at 1:200 for 5 days. Brains were then washed in PBS, dehydrated in an ascending ethanol series (30% to 100%), cleared and finally mounted in methyl salicylate (M6752, Sigma-Aldrich, France) for observation. Brains were scanned using a laser-scanning confocal microscope (Zeiss LSM 700) with a W Plan-Apochromat 20×/NA 1.0 objective using sequential excitation wavelengths of 488 nm and 555 nm, observing via two color-filtered channels around 510 nm and 590 nm, respectively.

### Honey bee preparation for *in vivo* calcium imaging

Transgenic workers were collected at emergence and were maintained in small cages in the dark at 34°C for 2 weeks. On the day of the experiment, they were individually chilled on ice for 5 min until they stopped moving. Then, they were prepared following the standard preparation used to image the ventral part of the honey bee brain [19]. Briefly, the honey bee was fixed in a plastic chamber with its antennae oriented to the front and the proboscis was fixed using beeswax to avoid movement of the brain during the experiment. A pool was built with beeswax and pieces of plastic around the head capsule, and a small window was then cut in the head cuticle. Glands as well as trachea were removed to expose the brain, and the pool was filled with ringer solution (in mM: NaCl, 130; KCl, 6; MgCl_2_, 4; CaCl_2_, 5; sucrose, 160; glucose, 25; Hepes, 10; pH 6.7, 500 mOsmol; all chemicals from Sigma-Aldrich, France), to avoid desiccation of the brain surface. The honey bee was then left 30 min in a moisturized and dark place before the calcium imaging.

### Calcium imaging

A T.I.L.L. Photonics imaging system (Martinsried, Germany) was used to perform *in vivo* optical recordings, as described elsewhere [19, 73, 74]. An epifluorescence microscope (Olympus BX51WI) was used to record activity in the different regions of the brain with either a 4× dry objective (Olympus, PlanCN; NA 0.10) for whole-brain recordings, a 10× water-immersion objective (Olympus, UMPlanFL; NA 0.3) for AL and MB recordings or a 20× water-immersion objective (Olympus, UMPlanFL; NA 0.5) for LH recordings. GCaMP6f was excited using 475 nm monochromatic light (T.I.L.L. Polychrom IV). The fluorescence signal was separated by a 505 nm dichroic filter and a long-pass 515 nm emission filter and recorded with a 640 × 480 pixels 12-bit monochrome CCD camera (T.I.L.L. Imago) cooled to −12°C with 4 × 4 binning on chip. Each measurement consisted of 100 frames recorded at a rate of 5 Hz (integration time for each frame ^~^50ms).

### Odor stimuli

A constant airstream was directed from a distance of 1 cm to the bee’s antennae, and odor stimuli were given at the 15^th^ frame for 1 s. For each odor stimulus (all obtained from Sigma-Aldrich, France), 5 μL of the solution were deposited on a filter paper inserted in a Pasteur pipette. A pipette containing a clean piece of filter paper was used as control stimulus.

For whole-brain recordings, we tested a small set of 3 odorant stimuli known from previous work to trigger strong neural activity: 2-heptanone, octanal, and heptanal. For AL and LH recordings, we tested 16 aliphatic odorants previously used in calcium imaging studies [19, 45] belonging to 4 functional groups (primary and secondary alcohols, aldehydes, and ketones) and having 4 different carbon chain lengths (6, 7, 8, and 9 carbons). The use of these odorants which are part of floral blends honey bees encounter while foraging [75] allowed to compare the results obtained in this study using GCaMP6f imaging with previous studies using classical calcium-sensitive dyes such as Fura-2 dextran [19] or Calcium green 2-AM [45]. These 16 odorants were used to record neural activity at the level of the AL and at the level of the LH on two different groups of bees. For recordings at the level of the MB, a reduced list of odorants which give strong neural activity was used: 1-hexanol, 2-hexanone, 2-heptanone, heptanal, octanal, and the air control. For LH recordings in responses to social pheromones, we tested both volatile (IPA and ocimene) and non-volatile compounds from the queen pheromone (QMP, 9-ODA, 9-HDA, HOB and HVA) and from the brood pheromone (methyl oleate, ethyl oleate, methyl linoleate, methyl palmitate, ethyl palmitate) at a concentration of 50μg/μL. As control stimulus, a pipette containing the solvent (isopropanol) or a clean piece of filter paper was used.

Each odorant stimulus was presented twice in a pseudo-randomized order, avoiding the consecutive presentation of stimuli with the same functional group or the same carbon chain length. Only animals in which all odorants in the panel were presented were kept for analysis.

### Data processing and analyses

All analyses were carried out using custom-made software written in IDL 6.0 (Research Systems, Boulder, CO). Each odor response signal corresponds to a three-dimensional array consisting of two spatial dimensions (*x*- and *y*-coordinates) along time (100 frames). First, the relative fluorescence changes were calculated as Δ*F*/*F* = (*F* – *F*_0_)/*F*_0_ by taking as reference background *F*_0_ the average of three frames just before the odorant stimulation (frames 9-11). Possible irregularities of lamp illumination and bleaching were corrected by subtracting the median pixel value of each frame from every single pixel of the corresponding frame. Finally, the two spatial dimensions were filtered with a Gaussian filter of window size 7×7 pixels for AL recordings or 3×3 pixels for LH and MB recordings for noise reduction. A biphasic calcium signal was observed in all recordings. As in previous studies using bath-applied Calcium Green [45], a high contrast measure for the intensity of the odor-induced response was obtained by averaging three consecutive frames at the end of the odor presentation (frames 19-21) and subtracting the average of 3 frames during the second, negative component of the signal (frames 49-51).

For the quantification of response intensity and similarity relationships, a mask was precisely drawn in order to exclude regions outside of the imaged structure. A pixel-wise analysis was then performed, using all the pixels within the mask. Response intensity was calculated by averaging the intensity of all pixels located within the unmasked area. Similarity relationships between neural activity patterns *I*(*x, y*) were calculated by measuring Euclidian distances

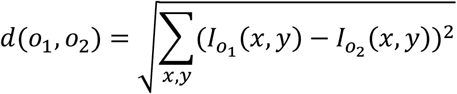

for all the 120 odorant pairs *o*_1_; *o*_2_ within each animal.

For all analyses, average values for the two presentations of each odorant were used except for the comparison of Euclidian distances for the same vs. different odorants (Fig 3E) in which each single odorant presentation was used.

We used the Euclidian distances between odorants calculated using behavioral experiments by Guerrieri *et al*. [48] to compare similarity relationships among odorants at the neural level with data obtained at the behavioral level. All results are displayed as means over individuals ± SEM.

### Statistical analyses

Odor response intensities were compared with ANOVA for repeated measurements, using odors as within-group factors. A Dunnett post hoc test was applied to compare the intensity of the response to each stimulus with a common reference, the air control. Responses to social pheromone compounds were compared using a Friedman ANOVA. Wilcoxon matched-pairs tests were applied to compare Euclidian distances between the same and different odors for AL recordings. Pearson correlation analyses were performed between response intensity and the logarithm of odorants’ vapor pressure in mmHg (16 aliphatic odors recorded in the AL and in the LH) and between response intensities recorded in this and previous studies (OSNs [45] or PNs [19]). A Fisher *z*-test was used to test for significant differences between linear regression following Pearson correlation (R package *multilevel, cordif* function). Mantel tests were used to compare matrices containing the Euclidian distances for the 120 possible odor-pair among 16 odorants, either between different imaging studies, or between neural and behavioral measures (R package *ade4, mantel.rtest* function). To explore similarity relationships among odorants, hierarchical clustering using Ward’s classification method was used (R package *stats*, *hclust* function). All tests were performed with GraphPadPrism V7.00 and R (www.r-project.org).

## ACKNOWLEDGEMENTS

We thank Robin Parlow, Eva Theilenberg, Marion Müller-Borg, and Virginie Larcher for technical assistance. We thank Sara Bariselli for her support with the egg injection and Amélie Cabirol for fruitful discussions. We also thank Maud Combe for developing the custom programs used for data analysis.

We thank the French Research National Agency (ANR to JCS, Project ANR-17-CE20-0003 Bee-O-Choc, Project ANR-22-CE37-0029 PHEROBRAIN), the CNRS (JCS) and the University Paris-Saclay (JC and JCS), the DFG (Deutsche Forschungsgemeinschaft to MB) and CIMeC, University of Trento (AH) for funding.

## AUTHOR CONTRIBUTIONS

J.C., M.O., A.H., J.C.S., and M.B. conceived the experiments. J.C. and M.O. collected the data. J.C., M.O., A.H., J.C.S., and M.B. analyzed the data and interpreted the results. J.C., A.H., J.C.S., and M.B. wrote the manuscript. All authors read and approved the final version of the manuscript.

## SUPPORTING INFORMATION

**S1 Fig.**
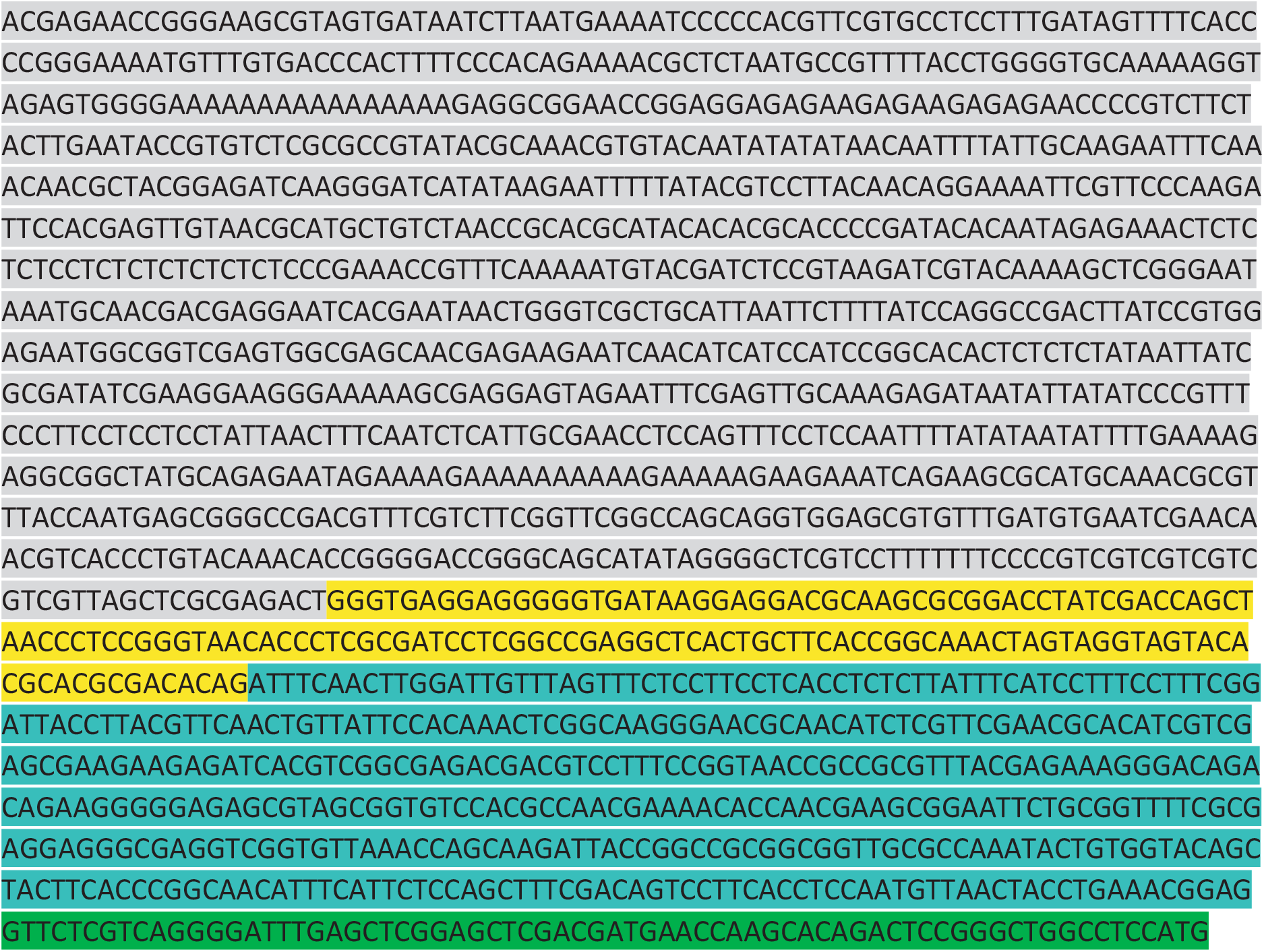
Synapsin promoter sequence of the honey bee. The upstream promoter region and the 5’ UTR sequence of the honey bee *synapsin* (*syn*) gene as revealed from the gene annotation release 104 (NCBI Apis mellifera Annotation Release 104; https://www.ncbi.nlm.nih.gov/genome/annotation_euk/Apis_mellifera/104/). In grey the predicted promoter and upstream region. Different colors indicate the different annotated exons that were fused to obtain a single 5’UTR region.

**S2 Fig.**
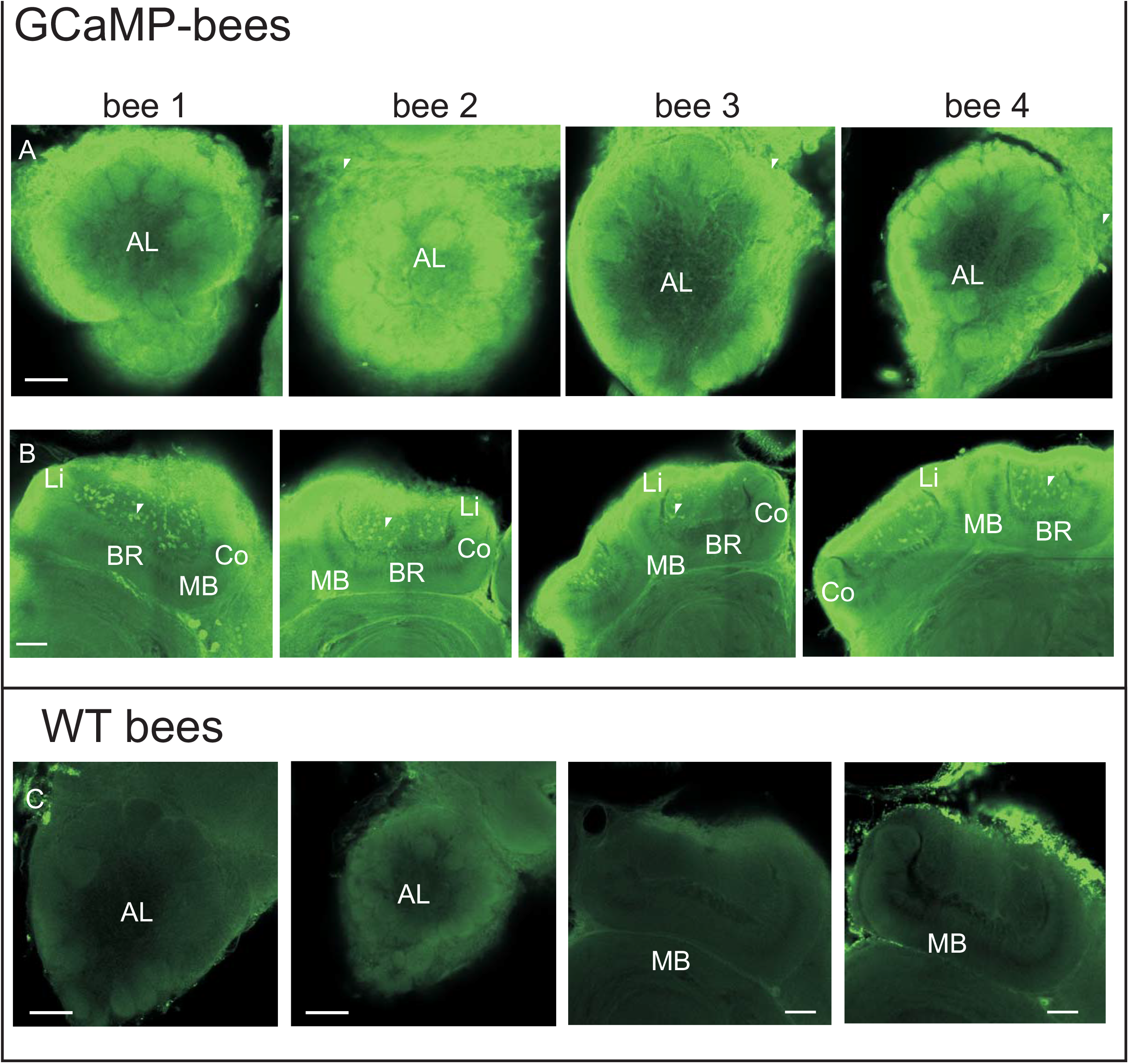
Comparison of immunostaining of GCaMP6f in different honey bees. **A-** GCaMP6f expression (anti-GFP in green) in the AL of different honey bees showing strong expression in the somata of projection neurons and local neurons (white arrows). **B** - GCaMP6f expression (anti-GFP in green) in the mushroom bodies in different honey bees showing strong expression in somata of the Kenyon cells (in the cup of the calyces, see white arrows). Scale bar = 50 μm. AL: Antennal lobe, MB: Mushroom body, Li: Lip, BR: Basal ring, Co: Collar. **C** - Examples of AL and MB of wild type (WT) bees showing no GCaMP6f expression.

**S3 Fig.**
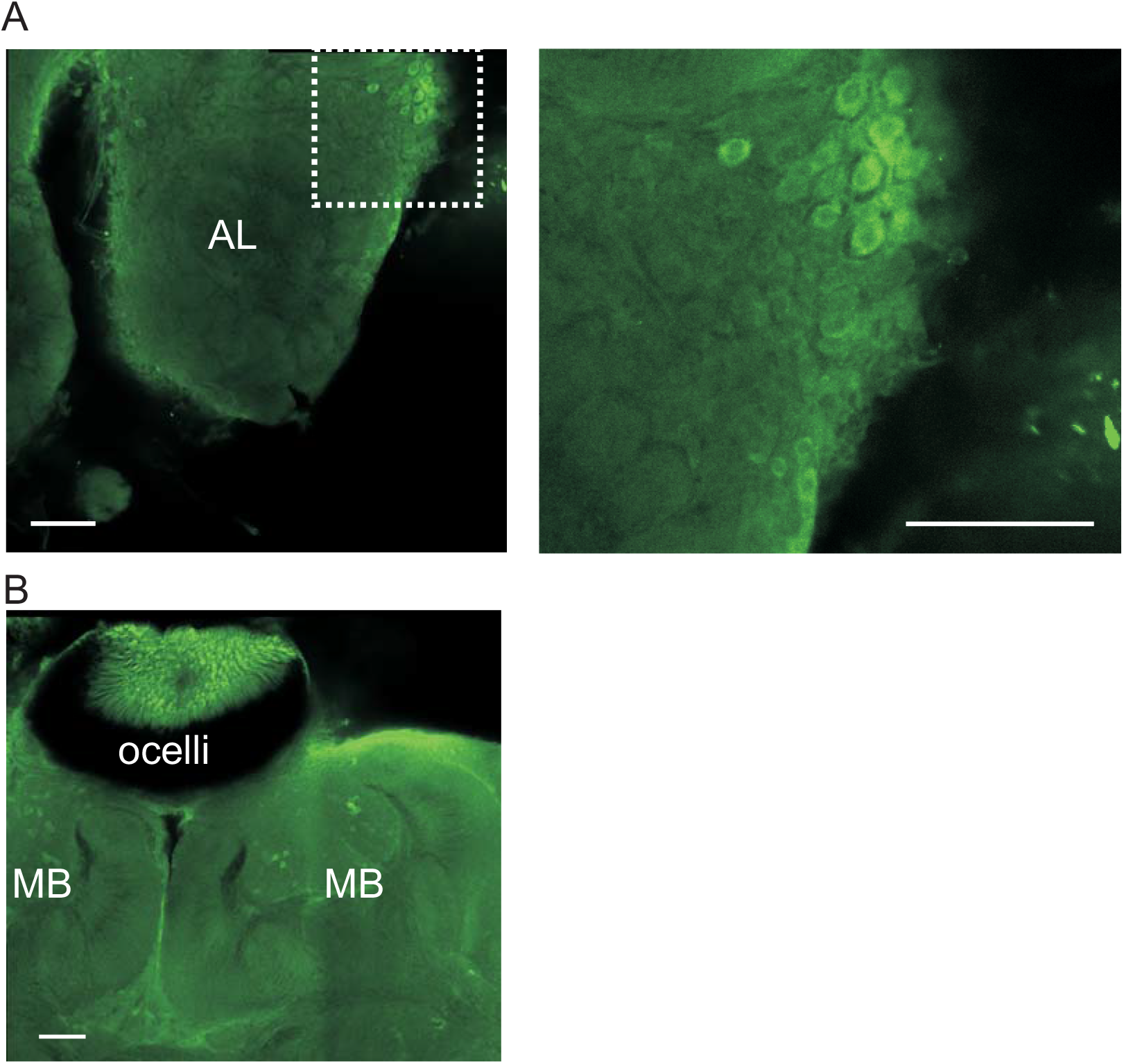
GCaMP6f expression in different regions. **A-** GCaMP6f expression (anti-GFP in green) in the lateral cluster of local/projection neurons near the AL (left) and its zoom (right), showing very broad expression in almost all the somata. Scale bar = 50 μm. AL: Antennal lobe. **B** - GCaMP6f expression (anti-GFP in green) in the photoreceptors of the ocelli. MB: Mushroom body.

**S4 Fig.**
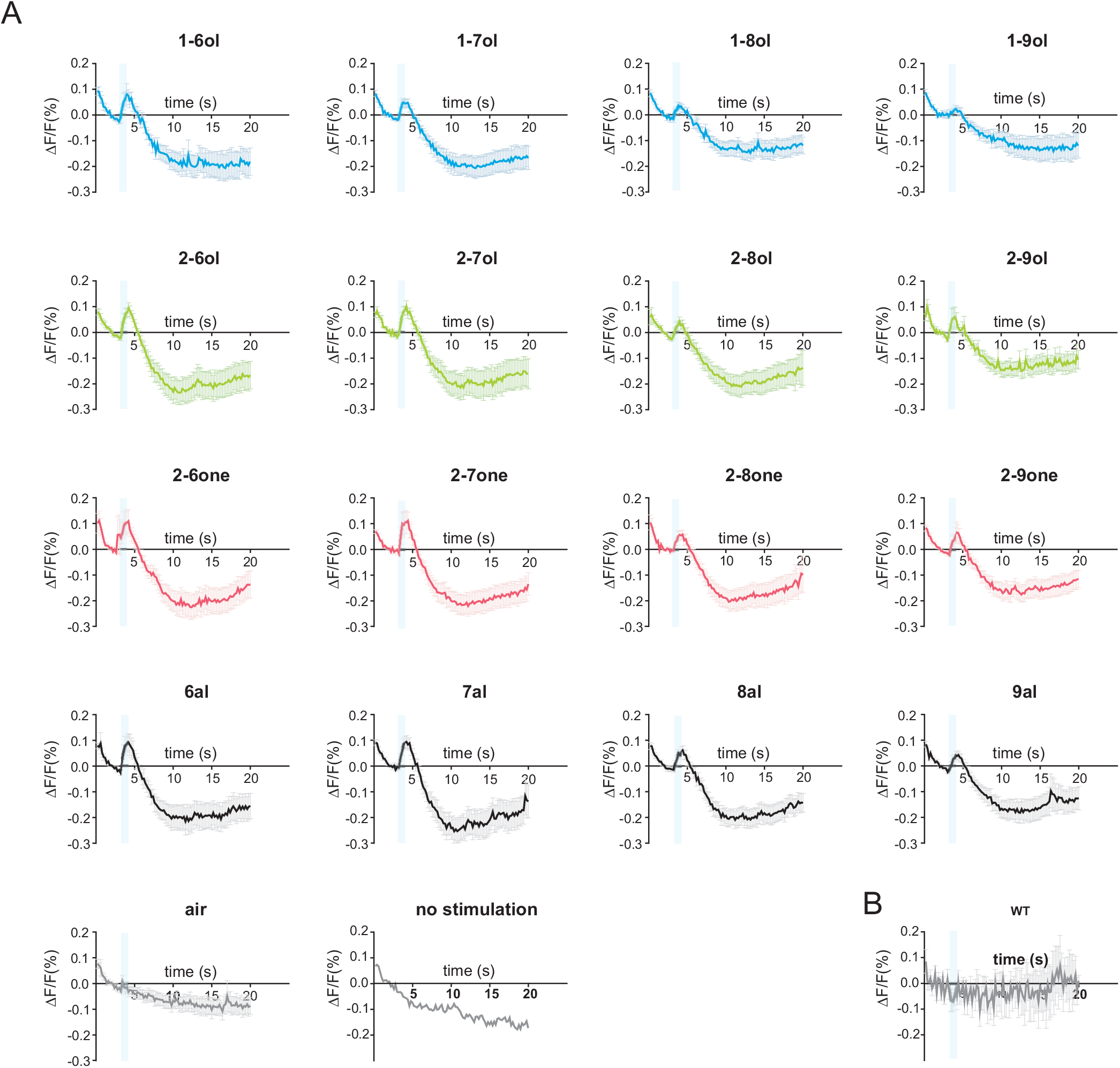
Time-courses of odor-evoked responses recorded in the antennal lobe. A- Time course of odor-evoked responses (Δ*F*/*F* [%]) recorded in the AL (*n* = 11 honey bees) to different odorants and the air control (in grey). Time course with no olfactory stimulation (*n* = 2 honey bees) is also shown (in grey). B- Time course (Δ*F*/*F* [%]) recorded in the AL of wild type bees (*n* = 10 honey bees) following odor presentations, showing no signal in non-transgenic bees. The data underlying the graphs shown in the figure can be found in S5 Data.

**S5 Fig.**
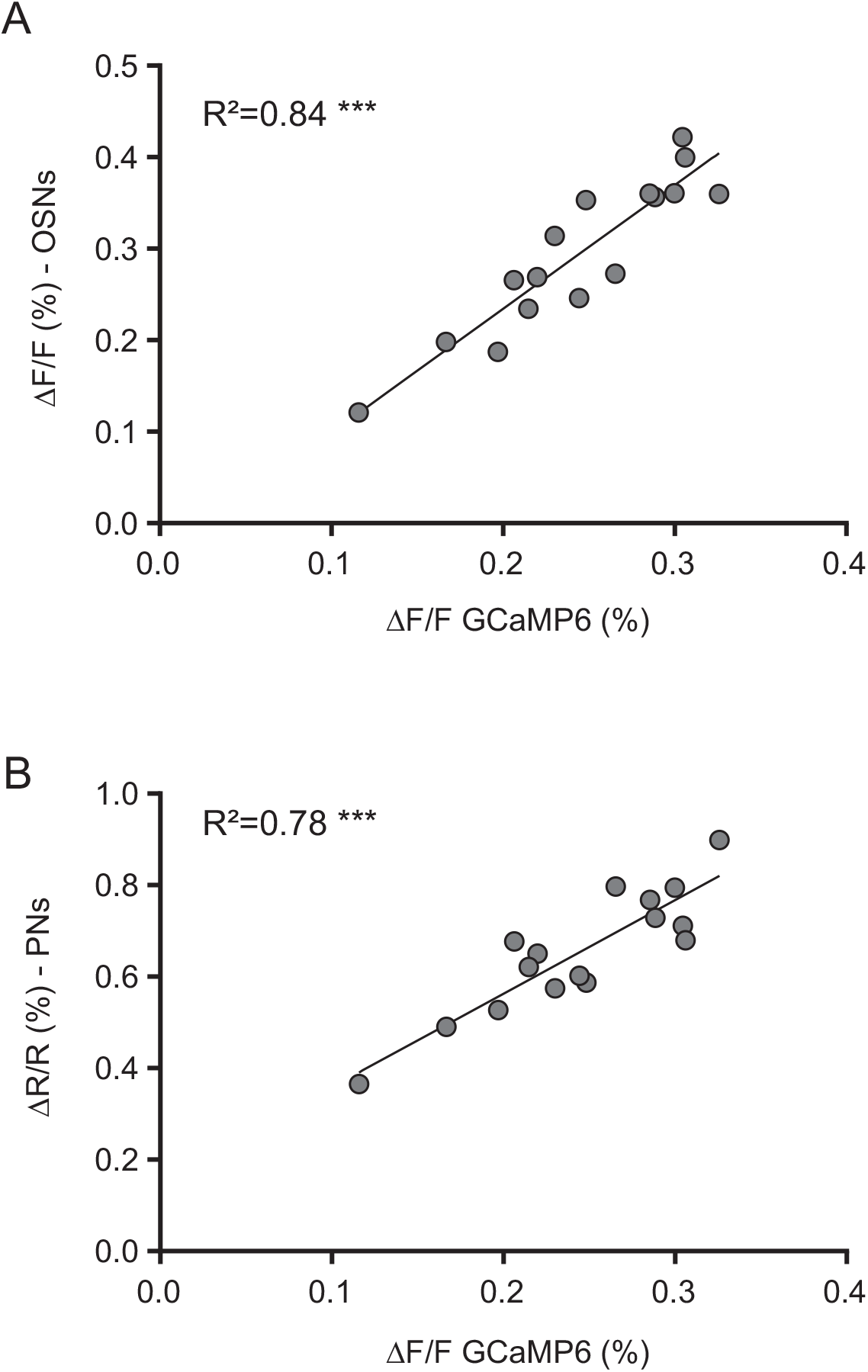
Correlation between amplitudes of calcium responses recorded in the antennal lobe. **A-** Amplitude of calcium responses (Δ*F*/*F* [%]) recorded in Olfactory sensory neurons (OSNs) in [45] as a function of the amplitude of calcium responses (Δ*F*/*F* [%]) recorded in the AL in this study. The linear regression shows a significant correlation (*R*^2^ = 0.84, *** *p* = 6e-7). **B-** Amplitude of calcium responses (Δ*F*/*F* [%]) recorded in Projection neurons (PNs) in [19] as a function of the amplitude of calcium responses (Δ*F*/*F* [%]) recorded in the AL in this study. The linear regression shows a significant correlation *R* = 0.78, *** *p* = 6e-6).

**S6 Fig.**
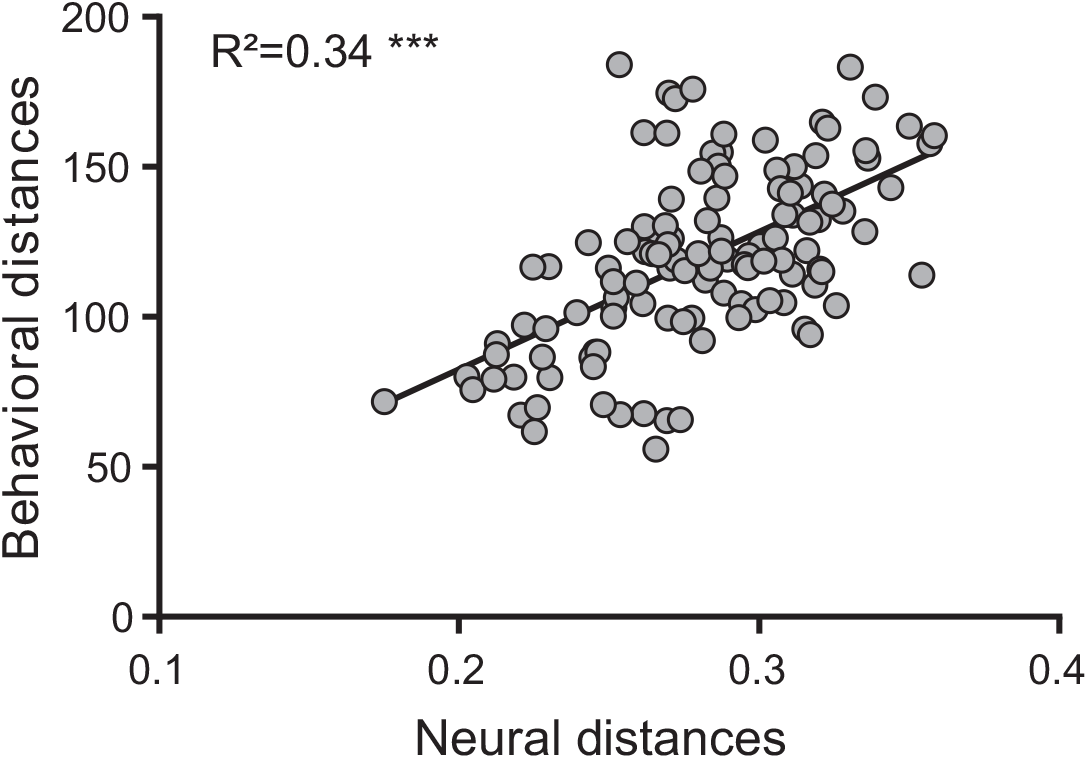
Correlation between neural and behavioral distances. Correlation of Euclidian distances recorded in the AL in this study are highly correlated with behavioral distances recorded in (48) *R* = 0.34, *** *p* < 0.001). The data underlying the graphs shown in the figure can be found in S8 Data.

**S7 Fig.**
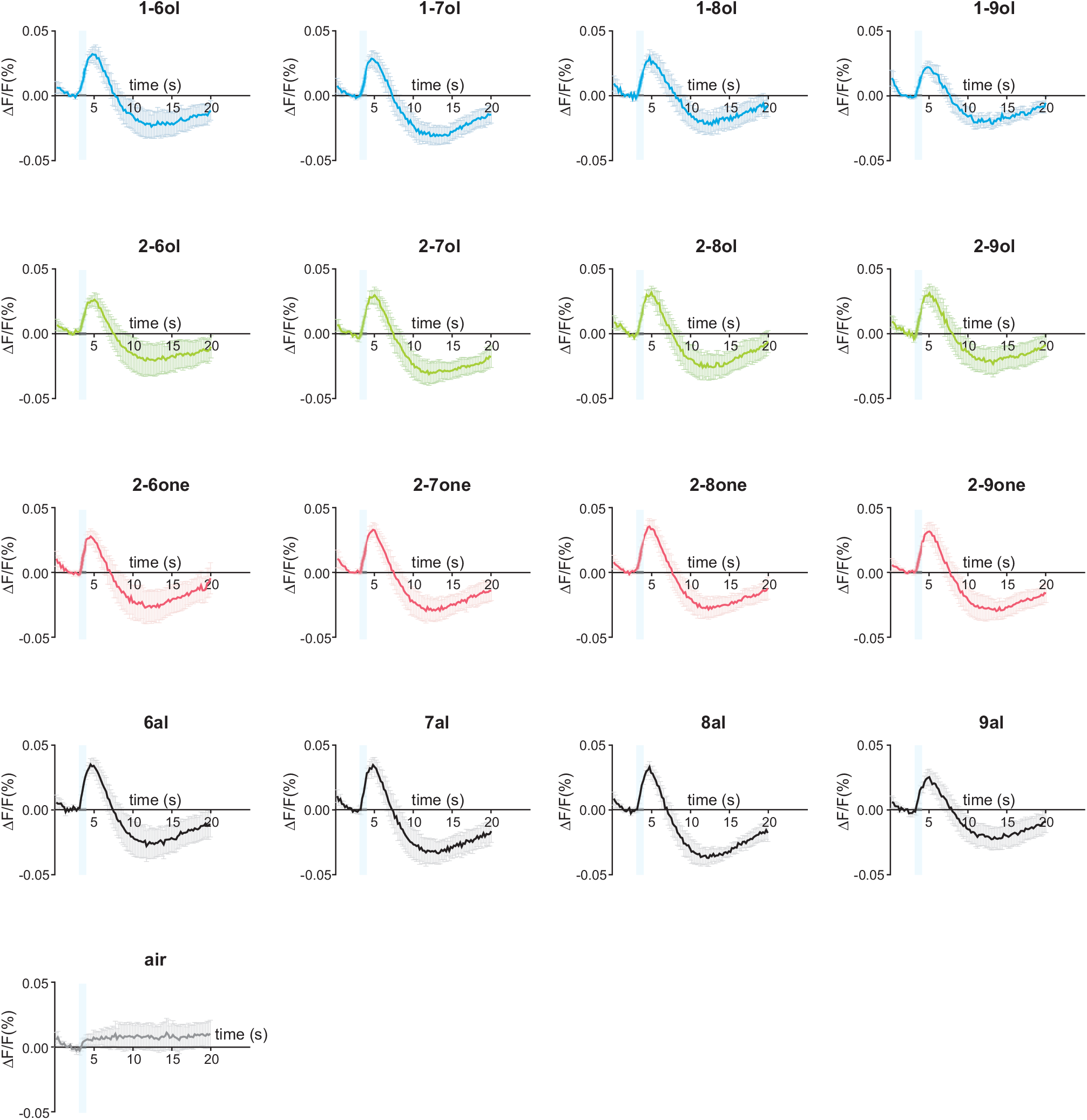
Time-courses of odor-evoked responses recorded in the lateral horn. Time course of odor-evoked responses (Δ*F*/*F* [%]) recorded in the LH (*n* = 8 honey bees) to the different odorants and to the air control (in grey). The data underlying the graphs shown in the figure can be found in S6 Data.

**S8 Fig.**
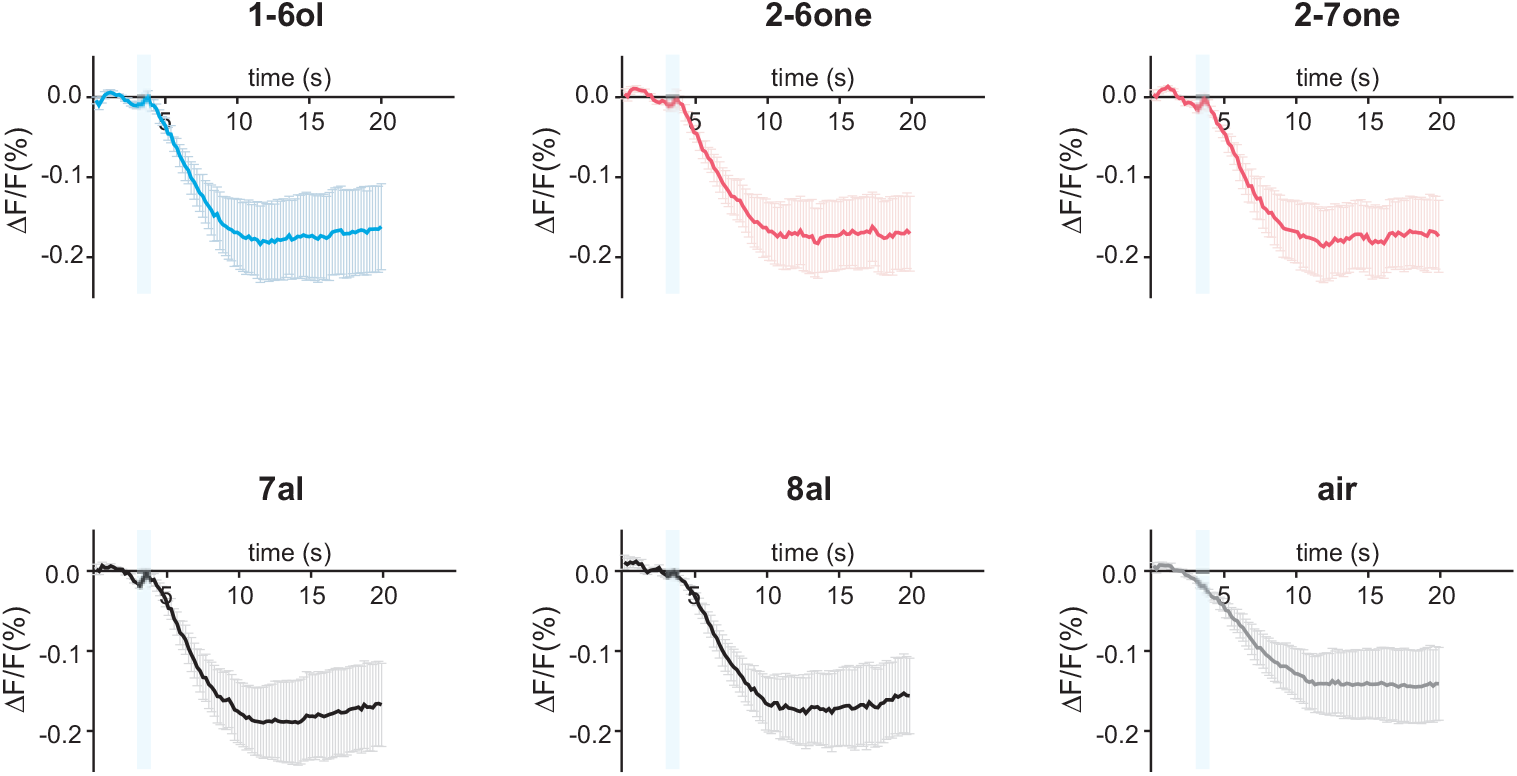
Time-courses of odor-evoked responses recorded in the mushroom bodies. Time course of odor-evoked responses (Δ*F*/*F* [%]) recorded in the MB (*n* = 6 honey bees) to the different odorants and to the air control (in grey). The data underlying the graphs shown in the figure can be found in S7 Data.

S1 Data. Calcium imaging data recorded in the antennal lobe. Data underlying Fig 3.

S2 Data. Calcium imaging data recorded in the lateral horn. Data underlying Fig 4.

S3 Data. Calcium imaging data recorded in the mushroom bodies. Data underlying Fig 5.

S4 Data. Calcium imaging data recorded in the lateral horn in response to pheromonal compounds. Data underlying Fig 6.

S5 Data. Timecourses data recorded in the antennal lobe. Data underlying Fig 3B.

S6 Data. Timecourses data recorded in the lateral horn. Data underlying Fig 4B.

S7 Data. Timecourses data recorded in the mushroom bodies. Data underlying Fig 5B.

S8 Data. Behavioral Euclidian distances. From Guerrieri et al. 2005; Data underlying S6 Fig.

